# A Novel Method for Improving the Accuracy of MR-derived Patient-specific Vascular Models using X-ray Angiography

**DOI:** 10.1101/2021.12.22.472309

**Authors:** John D. Horn, Zbigniew Starosolski, Michael J. Johnson, Avner Meoded, Shaolie S. Hossain

**Affiliations:** Molecular Cardiology Research Laboratory, Texas Heart Institute, Houston, TX, USA; Department of Radiology, Texas Children’s Hospital, Houston, TX, USA; Department of Radiology, Baylor College of Medicine, Houston, TX, USA; Oden Institute for Computational Engineering and Sciences, University of Texas at Austin, Austin, TX, USA

## Abstract

MR imaging is a noninvasive imaging modality that is commonly used during clinical follow up and has been widely utilized to reconstruct realistic 3D vascular models for patient-specific analysis. In a recent work, we utilized patient-specific hemodynamic analysis of the circle of Willis to noninvasively assess stroke risk in pediatric Moyamoya disease (MMD)—a progressive steno-occlusive cerebrovascular disease that leads to recurrent stroke. The objective was to identify vascular regions with critically high wall shear rate (WSR), signifying elevated stroke risk. However, sources of error including insufficient resolution of MR images can negatively impact vascular model accuracy, especially in areas of severe pathological narrowing, and thus diminish clinical relevance of simulation results, as local hemodynamics are sensitive to vessel geometry. We have developed a novel method to improve the accuracy of MR-derived 3D vascular models utilizing 2D X-ray angiography (XA), which is considered the gold standard for clinically assessing vessel caliber. In this workflow, “virtual angiographies” (VA) of 3D MR-derived vascular models are conducted, producing 2D projections that are compared to corresponding XA images guiding the local adjustment of modeled vessels. This VA-comparison-adjustment loop is iterated until the two agree, as confirmed by an expert neuroradiologist. Using this method, we generated models of the circle of Willis of two patients with a history of unilateral stroke. Blood flow simulations were performed using a Navier-Stokes solver within an isogeoemtric analysis framework and WSR distributions were quantified. Results for one patient show as much as 45% underestimation of local WSR in the stenotic left anterior cerebral artery (LACA) and up to a 60% underestimation in the right anterior cerebral artery when using the initial MR-derived model compared to the XA-adjusted model, emphasizing the need for verifying improved accuracy of the adjusted model. To that end, vessel cross-sectional areas of the pre- and post-adjustment models were compared to those seen in 3D CTA images of the same patient. CTA has superior resolution and signal to noise ratio compared to MR imaging but is not commonly used in clinic due to radiation exposure concerns, especially in pediatric patients. While the vessels in the initial model had normalized root mean squared deviations (NRMSDs) ranging from 26% to 182% and 31% to 69% in two patients with respect to CTA, the adjusted vessel NRMSDs were comparatively smaller (32% to 53% and 11% to 42%). In the mildly stenotic LACA of patient 1, the NRMSDs for the pre- and post-adjusted models were 49% and 32%, respectively. These findings suggest that our XA-based adjustment method can considerably improve the accuracy of vascular models, and thus, stroke-risk prediction. An accurate individualized assessment of stroke risk would be of substantial clinical benefit because it would help guide the timing of preventative surgical interventions in pediatric MMD patients.

## 1. Introduction

Patient-specific modeling of vascular networks has proven to be a valuable tool in the research of vascular pathologies and their treatment^1^. Researchers have used patient-specific vascular modeling to detect coronary artery disease^2^, to inform treatment planning via optimization of graft placement^3^, surgical intervention^4^ and nanoparticulate drug delivery^5, 6^, and to assess aneurysms and their treatments^7^. In recent work^8^, we have used patient-specific modeling to evaluate the hemodynamics within the circle of Willis (CoW) of pediatric patients with Moyamoya disease (MMD) – a progressive steno-occlusive cerebrovascular disease that can result in recurring stroke events^9^. Because there are no known treatments to reverse or slow the vessel narrowing caused by MMD, current clinical interventions focus on addressing the complications of the disease (e.g., recurrent stroke, brain volume loss). These include neurosurgical strategies that allow blood flow to bypass areas of stenosis thereby reducing stroke risk. Therefore, an individualized assessment of stroke risk, as a means of guiding the timing of preventative surgical interventions, would be of great clinical benefit. It has been postulated that local wall shear rate (WSR) in the CoW arteries may be an indicator for disease progression and elevated stroke risk in pediatric MMD patients^8^. Critically high local WSR values above a 5,000 s^-1^ limit could result in thrombus formation leading to ischemic stroke^10, 11^. Thus, by assessing WSR, stroke risk in MMD patients can be evaluated, and clinicians can plan individualized patient follow-up and surgical strategies.

Computational fluid dynamics (CFD) simulation has been successfully used to evaluate hemodynamic quantities in the major arteries of the CoW^8, 12, 13^. As variations in vessel architecture can strongly influence local hemodynamics including WSR^14^, it is critically important that computational modeling accurately captures a patient’s vascular anatomy. Computed tomography angiography (CTA) is often utilized to create three-dimensional (3D) patient-specific vascular models for CFD studies, and many segmentation techniques have been developed to improve model accuracy^15, 16^. While CTA imaging is generally desirable for this purpose, it is not commonly collected during MMD patient follow up due to radiation exposure concerns, particularly in pediatric patients. MR time of flight (MR TOF) data is an alternative for reconstructing 3D patient-specific models that is more commonly collected during clinical follow up of young MMD patients. Given tradeoffs between resolution and imaging time – an important factor since pediatric patients must be sedated during image collection – MR TOF data often has coarse voxel resolution. This can affect geometric accuracy, particularly in pediatric patient models, where some vessels of interest may only be one or two voxel widths in diameter (vessel diameters as small as 1 mm vs. voxel widths near 0.5 mm). Further, important vessel features, such as severe MMD-caused vessel narrowing, can sometimes go unresolved by MR TOF which produces poor signal in areas of slow blood flow^17^. Additionally, vessels oriented parallel to the imaging slice plane can be inadequately resolved. Therefore, the accuracy of MR-derived models may not be adequate to reliably predict stroke risk in pediatric MMD patients. X-ray angiography (XA) is also often used to visualize a patient’s vasculature during clinical follow up to assess degree of pathological vessel narrowing. While these 2D projections obtained from XA cannot be used alone for 3D geometry reconstruction, they offer accurate depictions of vessel diameter and MMD-related vessel narrowing is typically well resolved. Lumen boundaries are also generally better defined in 2D XA images than 3D MR or CTA images.

Given the need for accurate vascular models for stroke risk assessment in MMD patients, as well as for other research applications in related fields, we have developed a method of 3D vascular model reconstruction that uses a combination of MR and XA patient imaging data. We hypothesize that, by using XA images as a reference for correct vessel caliber, MR-derived 3D models can be adjusted to improve accuracy. Our method involves computing 2D projections of the MR-derived 3D models onto a virtual detector plane using geometric parameters that mimic the clinical XA imaging setup (i.e., patient position in relation to the x-ray source and detector). The 2D projections are compared to the XA images and vessels in the 3D model are locally adjusted until the two agree. The effectiveness of the proposed adjustment methods in improving geometric accuracy is evaluated by comparing the resulting CoW models to 3D CTA images that were also available in the patient records. Treating the CTA images as an accurate 3D representation of the patient’s vascular anatomy, we can then assess if the MR-derived models have improved accuracy after XA-based adjustment.

## 2. Methods

In this work, we generated CoW models from de-identified imaging data of two patients who were retrospectively selected from a Texas Children’s Hospital database under an institutional review board– granted waiver of written authorization for consent. The selection criteria were 1) the availability of MR TOF, XA, and CTA imaging data in patient history, and 2) a history of stroke with visible vascular narrowing in the affected side. Patient 1 is diagnosed with MMD and presented with a stroke involving the right middle cerebral artery (RMCA). Patient 2 is a non-MMD patient, who suffered a right-side stroke. Evidence of severe occlusion in the RMCA was found on imaging. While unresolved, dehydration, inflammation, and viral infection including possible COVID infection, were suggested as potential causes for the stroke in patient 2.

### 2.1. Vascular model creation

To create patient-specific models of the CoW suitable for blood flow simulations we employ a four-stage process whereby an initial representation is segmented from clinical MR TOF patient data and adjusted using clinical XA imaging data. This model creation process is summarized in Figs. 1 and 2 and detailed in the following sections.

**Fig 1:**
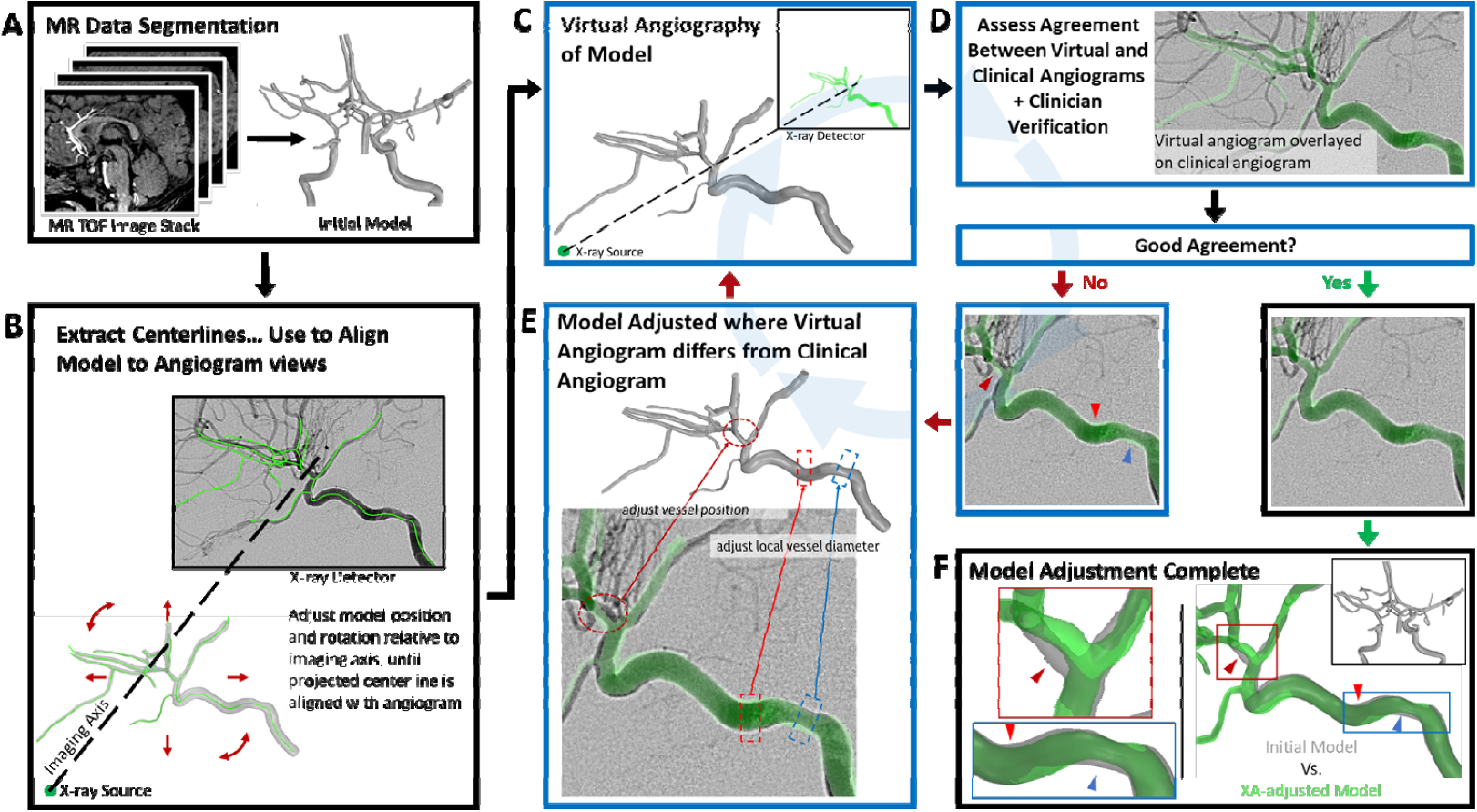
Graphical flow chart showing processes for model creation and XA-based adjustment. (A) An initial model is created by segmenting an MR TOF image stack. (B) Model centerlines are used to align the model relative to a virtual x-ray source and detector mimicking clinical XA image acquisition. (C) A virtual angiogram is computed of the model and (D) compared to the corresponding clinical XA image. If the virtual angiogram of the model and the clinical angiogram differ, the model is locally adjusted (E), and a new virtual angiogram is computed. The adjust-virtual angiography-compare steps (C-D-E) are iteratively repeated until there is good agreement with between the two angiograms. (F) Comparison of final adjusted model to initial model.

**Fig 2:**
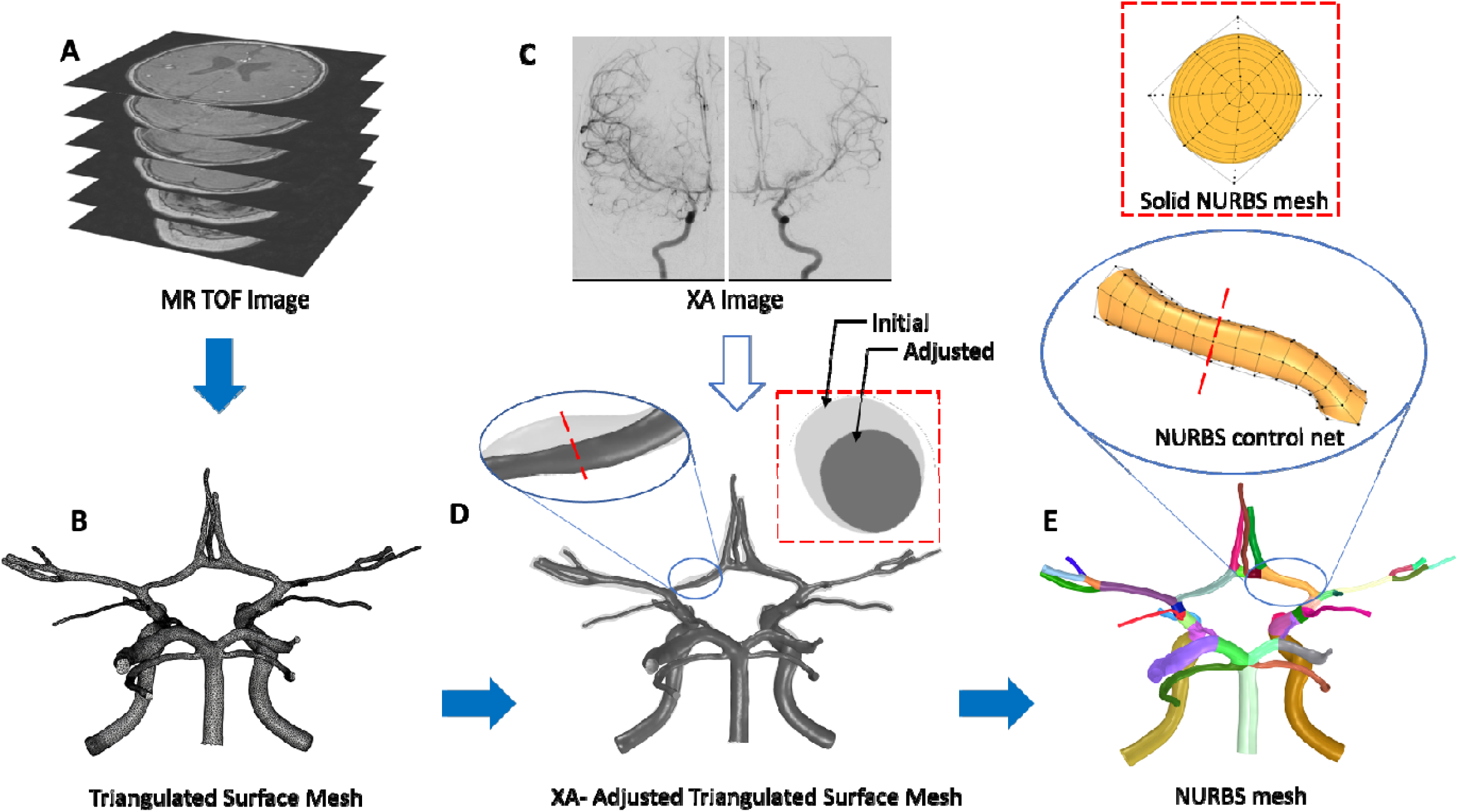
The vascular modeling pipeline. The Circle of Willis vasculature is segmented from MR TOF images (A) as a triangulated surface mesh (B) using 3D Slicer. 2D projections of the segmented mesh are registered to (C) XA images. The vessel diameters along the mesh are locally adjusted to match XA images. (D) shows the adjustments made across the entire mesh. Adjustments made along the LACA are highlighted in the insets. The adjusted surface mesh is used to generate the NURBS mesh (E). Each vessel corresponds to a volumetric NURBS patch. The insets show the NURBS patch, control net, and a cross section for the RACA.

#### 2.1.1. Generating the initial 3D model

We start with MR TOF data output as an image stack in a .NRRD file format. Prior to segmenting the CoW from the image stack, 3D Slicer (version 4.10.2) is used to resample the volume to yield voxel sizes with linear dimensions reduced by at least one half. This improves the segmentation results, especially in smaller vessels that are at most one or two voxels in diameter prior to the resampling. 3D Slicer is then used to segment the resampled image stack using intensity thresholding and other manual methods. The segmented label map is then exported as a triangulated surface mesh as a .STL file. The surface mesh is then post-processed using MeshMixer (version 3.5, Autodesk Inc., San Rafael CA) to locally smooth the segmented surface and repair any mesh errors resulting from the segmentation process. The resulting triangulated surface mesh is exported as a new .STL file and is considered the initial model or mesh (Fig. 1A).

#### 2.1.2. Aligning the 3D model with XA imaging axis

To compare vessel geometry of the CoW model to that seen on XA images, we must align the model to the XA imaging view. Metadata from XA imaging including *distance source to patient, distance source to detector, positioner primary angle*, and *positioner secondary angle* are used to define the imaging axis in a virtual space, mimicking the XA setup for each XA image. Centerlines, extracted from the CoW model, are used to find a position and orientation, relative to the imaging axis and virtual x-ray source, such that the CoW model centerlines align with the XA image when projected along the imaging axis onto the virtual detector plane (Fig. 1B). This is done by selecting a set of reference points on model the centerlines at vessel intersections and target points on the XA images that correspond to the same vessel intersections. Using custom algorithms implemented in Matlab (version 2019b, The MathWorks Inc, Natick MA), a range of a rotations and translations about the three coordinate axes are then tested to determine which model orientation and position results in the least average distance between the projected reference points and the target points. Using this process, the model is aligned to each of the XA images.

#### 2.1.3. Algorithmic adjustment of 3D model geometry using XA images

After determining the alignment of the initial model for each XA image, Rhino and Grasshopper (version 6, Robert McNeel & Associates, Seattle WA) are used to apply algorithmic adjustment to the CoW model based on each XA image. In this process, local vessel diameters of the CoW model are measured at set intervals along each vessel. After superimposing the clinical XA image on the virtual detector plane, vessel lumen edges are manually delineated on the XA images and confirmed by an expert neuroradiologist, and target vessel diameters are determined. The model diameters are then projected onto the virtual detector plane (Fig. 1C) and compared to the target diameters to generate scale factors that are used to locally expand or contract the 3D model radially about the vessel centerline (Fig. 1D). This process is repeated for each XA image. After adjustment in all six views—that is coronal and sagittal views of both the left (L) and right (R) internal carotid arteries (ICAs), and coronal and sagittal views of the basilar artery (BA)— the mesh is manually adjusted using Meshmixer to smooth out any unwanted artifacts from the algorithmic adjustment and to manually adjust at vessel intersections where the algorithmic process cannot be applied.

#### 2.1.4. Manual adjustment of 3D model geometry to achieve agreement with XA images

Finally, a series of manual adjustments are made to the 3D model to improve its agreement with the XA images. Virtual angiographies of the adjusted model onto the detector plane of each XA image are computed after orientating the model to the imaging axis based on the alignments determined earlier. The virtual angiographies result in 2D projections of the model which are overlaid onto the XA images enabling comparison of the model’s vessel geometry to the vessel geometry shown on the XA images. The model is manually adjusted using Meshmixer in areas where the projection does not agree with the vessel geometry as depicted on the XA image (Fig. 1E). This manual adjustment-projection-compare process is iteratively repeated until good agreement is achieved, as confirmed by an expert neuroradiologists. The final adjusted model (Fig. 1F) is then exported as a surface mesh and processed for nonuniform rational B-spline (NURBS) mesh generation.

#### 2.1.5 Reconstruction of solid NURBS model

The adjusted surface mesh from the previous step is used to generate a volumetric nonuniform rational B-spline (NURBS) reconstruction of the CoW (Fig. 2). This is done according to the procedures defined in a previous report^18^. The NURBS generation uses a template-based vascular modeling^19^ software built with the computer-aided design (CAD) package Rhinoceros 3D. First, a skeletonization algorithm is applied to the surface mesh to extract the vessel centerlines and topology. Along each centerline, a set of minimal torsion perpendicular frames are constructed at regular intervals. At branch points, where multiple vessel centerlines intersect, the frames are folded to define conforming NURBS patch interfaces. The fold angle is interpolated along a length of the centerline to avoid self-intersection and provide a higher quality parameterization. A closed intersection curve between the mesh and each frame is computed and interpolated with a NURBS curve. The control points of the NURBS curves are then lofted together to create the control net for the multi-patch NURBS surface reconstruction of the adjusted surface mesh. Finally, the surface control points are extruded in the radial direction to the centerline to obtain a volumetric NURBS reconstruction of the CoW that is used for analysis. Element sizes and refinements are chosen according to mesh independence studies^8^. Each branch in the vasculature is a quadratic NURBS solid with 16 elements in the circumferential direction and 17 elements in the radial direction. The element size in the radial direction is graded such that there are more elements near the boundary to resolve the boundary layer in the blood flow simulation. The number of elements in the axial direction depend on the length of branch with a mean element size of 0.75 mm. The NURBS geometry is maximally smooth in the radial and axial directions and contains four ***C*°** knot lines in the circumferential direction corresponding to the folding axes.

### 2.2. Blood flow simulations

Fig. 3 illustrates the simulation set up. The details of the governing equations and solution strategy can be found in previous works^8^ and references therein. Briefly, blood flow is assumed to be governed by the unsteady Navier-Stokes equations subjected to boundary conditions as follows. A pulsatile inflow condition^20^ is imposed at the three inlets: LICA, RICA, and BA, with a parabolic inflow profile^21^. A no-slip Dirichlet boundary condition is set on the vessel wall, and a traction-free boundary condition is prescribed at all the outlets for simplicity^8^. Blood is assumed to be a Newtonian fluid with a density of 1060 kg/m^3^ and a dynamic viscosity of 0.0035 Pa-s. The system of equations is solved by implementing a residual-based variational multiscale method using a Newton–Raphson procedure with a multistage predictor–corrector algorithm applied at each time step. The generalized-alpha method is used for time advancement. Within an isogeometric analysis framework^22^, quadratic NURBS are used to describe both the geometry and the solution space. The CoW model of patient 1 and patient 2 comprised 138,720 and 149,600 volumetric quadratic NURBS elements and is divided into 40 and 34 subdomains, respectively, where each subdomain is assigned to a compute core. WSR is computed from the equation of wall shear stress vector: **τ** = **(σ·n)-((σ·n)·n)n**, where **(σ·n)** is the traction vector, **σ** is the stress tensor, and **n** is the unit normal.

**Fig 3:**
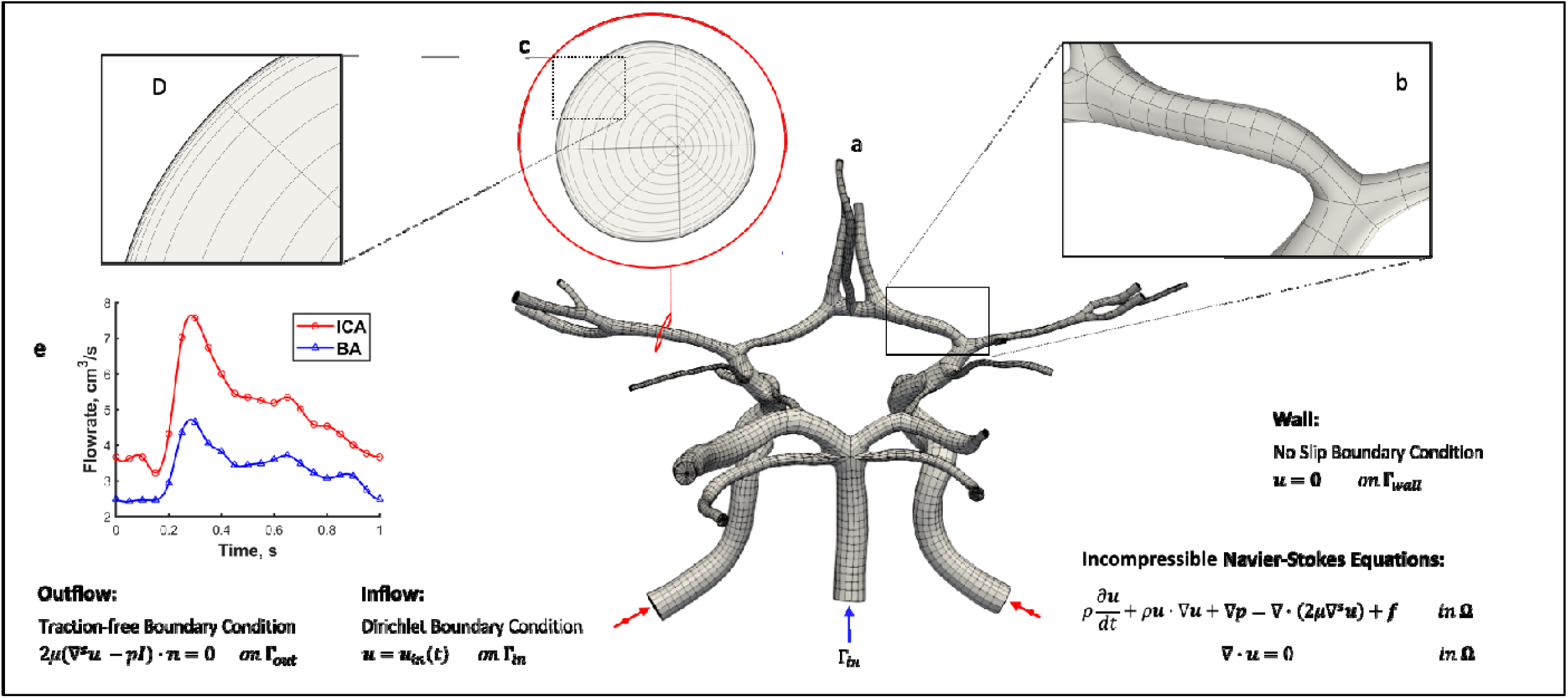
A volumetric NURBS mesh of the CoW (a) is generated from the adjusted CoW surface mesh. A close-up of a bifurcation (b) is shown to highlight the mesh quality and refinement used in the computation. The cross-section of the NURBS mesh (c) is shown to highlight the boundary layer refinement (d) used in the simulation. The equations and boundary conditions for the blood-flow simulation is presented, where u represents velocity, p is pressure, f is the external force, μ is dynamic viscosity, ρ is density, t is time, and n is the unit normal. A pulsatile inflow condition^20^ is prescribed at the three inlets. The inflow at the ICAs and BA are given by the red and blue waveforms respectively. A no-slip boundary condition is imposed along the vessel walls, and a traction-free boundary condition is applied at each outlet.

### 2.3. Verification of model accuracy

To determine if the XA-adjustment process described above improves accuracy of the reconstructed CoW geometry, we use CTA imaging data as a reference. For our analysis, the working assumption is that CTA images provide a more accurate 3D representation of vessel caliber, especially in areas of vessel narrowing compared to MR TOF. Thus, by comparing the MR-derived XA-adjusted models of a patient’s CoW to corresponding CTA imaging, we can determine whether the XA-adjusted model more closely represents the patient’s geometry than the MR-derived initial model. To do this, we quantify how much deviation exists between vessel cross-sectional areas taken from the models (initial and adjusted) and the corresponding vessel cross-sectional areas extracted from the CTA images. For our analysis, we focus on the major vessels of the anterior circulation: terminal ICAs, proximal middle cerebral arteries (MCAs), and proximal anterior cerebral arteries (ACAs)^23^.

#### 2.3.1. MR and CTA volume co-registration

First, we co-register the CTA image stack and the MR image stack from which the initial 3D model was segmented. This brings the vessels in the 3D model in approximate alignment with the CTA images and allows for direct comparison between the two. Prior to co-registration, regions of interest in each set of images were defined to isolate the vessels of interest in the CoW and reduce vessel misalignment related to high pixel intensities corresponding to boney structures. Rigid co-registration (translation and rotation only) was performed in 3D slicer using the built-in general registration module that optimizes alignment using a Mattes Mutual Information cost metric.

#### 2.3.2 Vessel cross sections

Vessel cross sections are obtained from the initial and adjusted models using planes perpendicular to the respective vessel centerlines (Fig.4a). These are taken at an interval of 0.1 mm along each centerline. These planes are then used to extract slices from the co-registered CTA volume resulting in perpendicular cross sections of each vessel that correspond to the model cross sections.

**Fig. 4:**
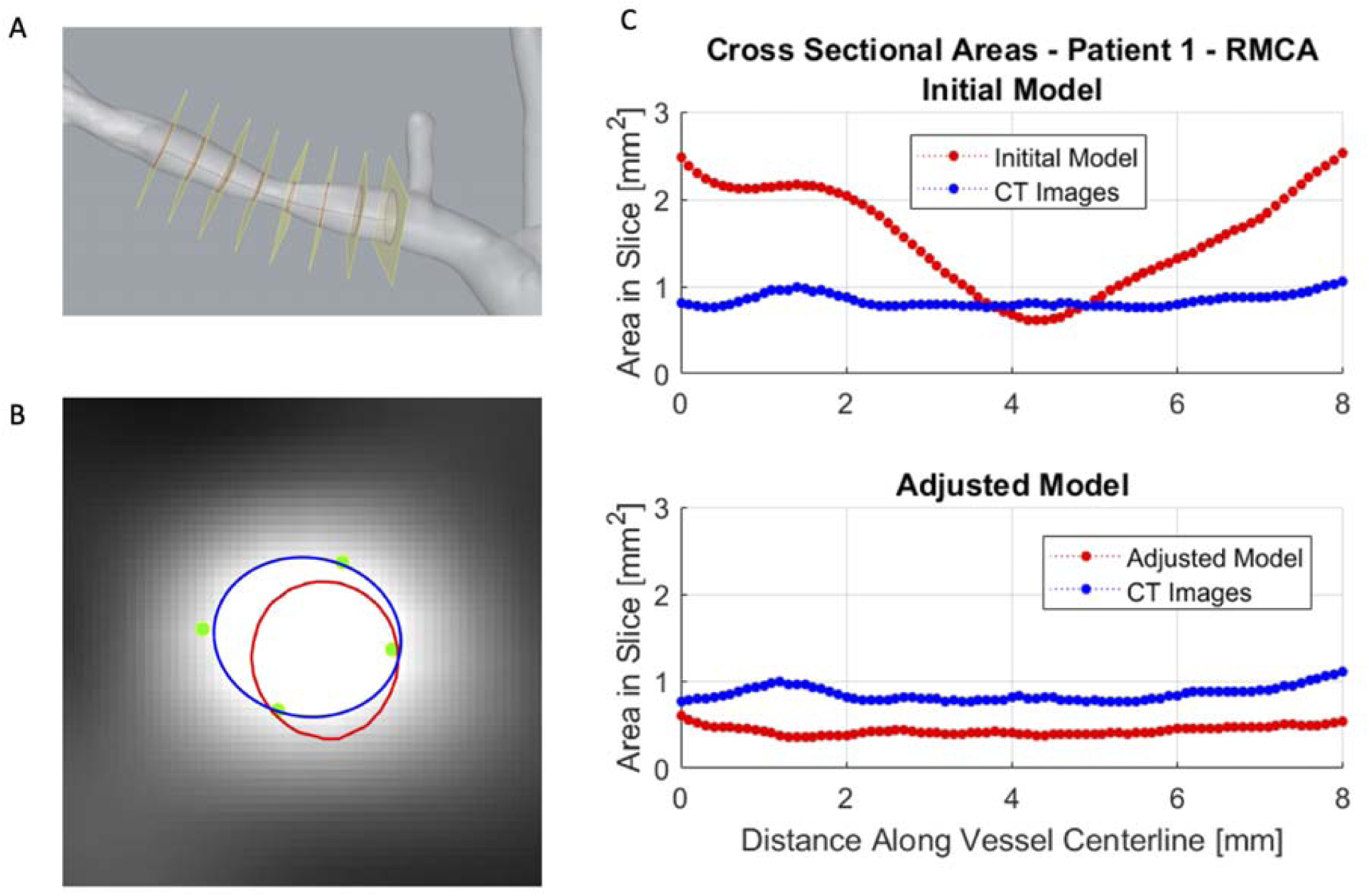
Overview of XA-adjustment verification strategy. (A) Several planes (yellow) along the length of a vessel of the 3D model (RMCA of the initial patient 1 model shown), perpendicular to its centerline, are used to extract vessel cross sections (red curves). (B) An example CTA slice corresponding to one of the perpendicular planes in (A). The cross section of the model superimposed on the CTA slice is shown by the red curve. Determination of the vessel lumen boundary on the CTA slice (blue curve) is guided by assessment of lumen extent provided by a trained neuroradiologist (green dots). (C) Cross-sectional areas of the model vessel and CTA vessel are computed in each slice and plotted against the position along the vessel centerline. The top plot compares the initial model (red curve) to the CTA-extracted lumen area (blue curve) and the bottom plot compares the adjusted model (red curve) to the CTA images (blue curve). These curves are used to compute the RMSD for each vessel.

#### 2.3.3. Detection of lumen boundary on CTA

With the goal of comparing cross-sectional lumen areas between the CoW model geometry and the CTA images, we extract the lumen boundary from each cross-sectional slice along the vessel centerline (Fig. 4a). The lumen boundary is detected based on an analysis of CTA pixel intensity taken along a line extending from the center of the vessel (greatest intensity) to beyond the lumen boundary (least intensity). The inflection point of this pixel intensity profile is identified along with the amplitude between the peak intensity and the lowest intensity. The lumen boundary is taken as the point where the pixel intensity is equal to the inflection point plus an offset value. This offset value, defined as a percentage of the profile amplitude, is calibrated using neuroradiologist input. This process is repeated at several angular positions relative to the center of the vessel producing a series of points outlining the lumen. To calibrate the offset value, a sample set of vessel cross sections was provided to an expert pediatric neuroradiologist who estimated vessel caliber from clinical CTA images. The neuroradiologist marked 3-5 points on each image corresponding to his visual estimate of the lumen boundary (Fig. 4b). Calibration of the offset value is achieved by determining the offset value that produces the lumen boundary with the least average distance to the points marked by the neuroradiologist. The optimal offset value is identified for each slice, and a mean offset for each vessel is calculated and used for detecting the lumen boundary.

#### 2.3.4. Comparison of cross-sectional areas

Cross-sectional areas of the initial and adjusted models are found by computing the areas enclosed by the rings produced by intersecting the perpendicular planes with the model surface meshes. Likewise, corresponding cross-sectional areas are computed using the detected lumen boundaries on CTA. As an example, Fig. 4c shows these cross-sectional areas for one vessel (patent 1 – RMCA) plotted against the position along the vessel centerline. To compare the cross-sectional areas of the models and to that of extracted from CTA images, along the length of each vessel, the root means squared deviation (RMSD) was computed,

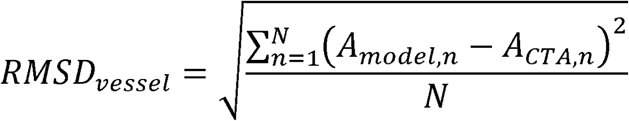

Where *A*_*model,n*_ and *A*_*CTA,n*_ are the cross-sectional areas in the n^th^ slice taken from the model and CTA images, respectively, and N is the total number of slices taken for that vessel. The RMSD values are then normalized by the mean cross-sectional area, taken from the CTA images, for a given vessel. The normalized RMSD (NRMSD) is then given by:

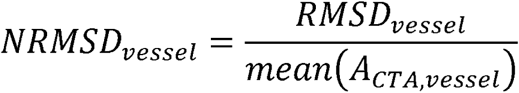

## 3. Results

Clinical imaging data, including XA and MR TOF, was used to create two patient-specific CoW models. The initial models, derived from the MR TOF volumetric data alone, were adjusted using XA images and the procedures outlined above. A comparison of the initial and adjusted models for the two patients can be seen in Fig. 5. The XA-based adjustment of patient 1’s CoW model generally resulted in vessels with reduced caliber. In contrast, the XA-based adjustment of the patient 2 model resulted in larger vessel calibers in general.

**Fig 5:**
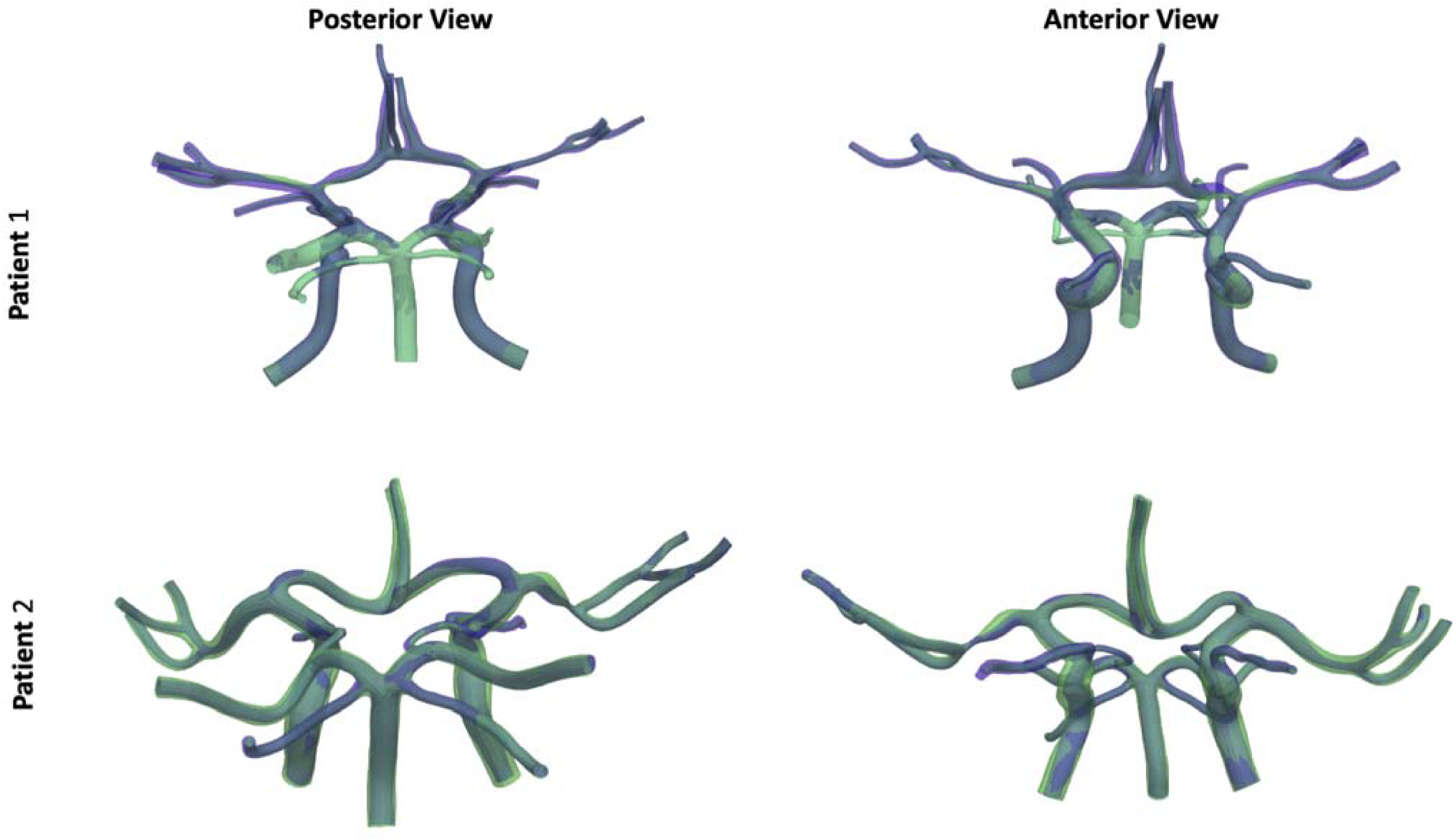
Comparison of initial (blue) and XA-adjusted models (green) for patient 1 (top panels) and patient 2 (bottom panels) shown in posterior (left) and anterior views (right).

In recent work^8^, we have used CoW models derived from pediatric MMD patient imaging data to assess stroke risk through blood flow simulation and an analysis of predicted WSR distributions. We conducted similar simulations using the initial and XA-adjusted CoW models for the two patients included in the present study and compared the WSR distributions of the pre- and post-adjustment models (see Fig. 6). For patient 1, the initial model resulted in a surface-averaged peak systolic WSR of 5,290 s^-1^ and the adjusted model produced a surface-averaged peak WSR of 9,830 s^-1^. The adjusted model produced focal areas with greater maximum WSR at peak systole than the initial model. Of note, the proximal left MCA (LMCA) and left ACA (LACA) of the adjusted model had a maximum WSR of 50,300 s^-1^ and 37,000 s^- 1^, whereas the corresponding vessels of the initial model had a maximum WSR of 37,800 s^-1^ and 20,400 s^-1^, respectively. On the other hand, in proximal right ACA (RACA) and right MCA (RMCA), maximum WSRs of 18,000 s^-1^ and 17,900 s^-1^ were predicted in the initial models, respectively, and a maximum WSRs of 44,900 s^-1^ and 38,400 s^-1^ were predicted in the adjusted models, respectively. For patient 2, the initial model resulted in a surface-averaged peak systolic WSR of 7,940 s^-1^ and the adjusted model produced a surface-averaged peak WSR of 4,763 s^-1^. In the initial model, the LMCA saw a peak WSR of 40,600 s^-1^, while the same vessel in the adjusted model saw a maximum WSR of 10,200 s^-1^. The RACA in the initial model had a peak WSR of 9,200 s^-1^, while the adjusted RACA had a peak WSR of 25,800 s^-1^.

**Fig 6.**
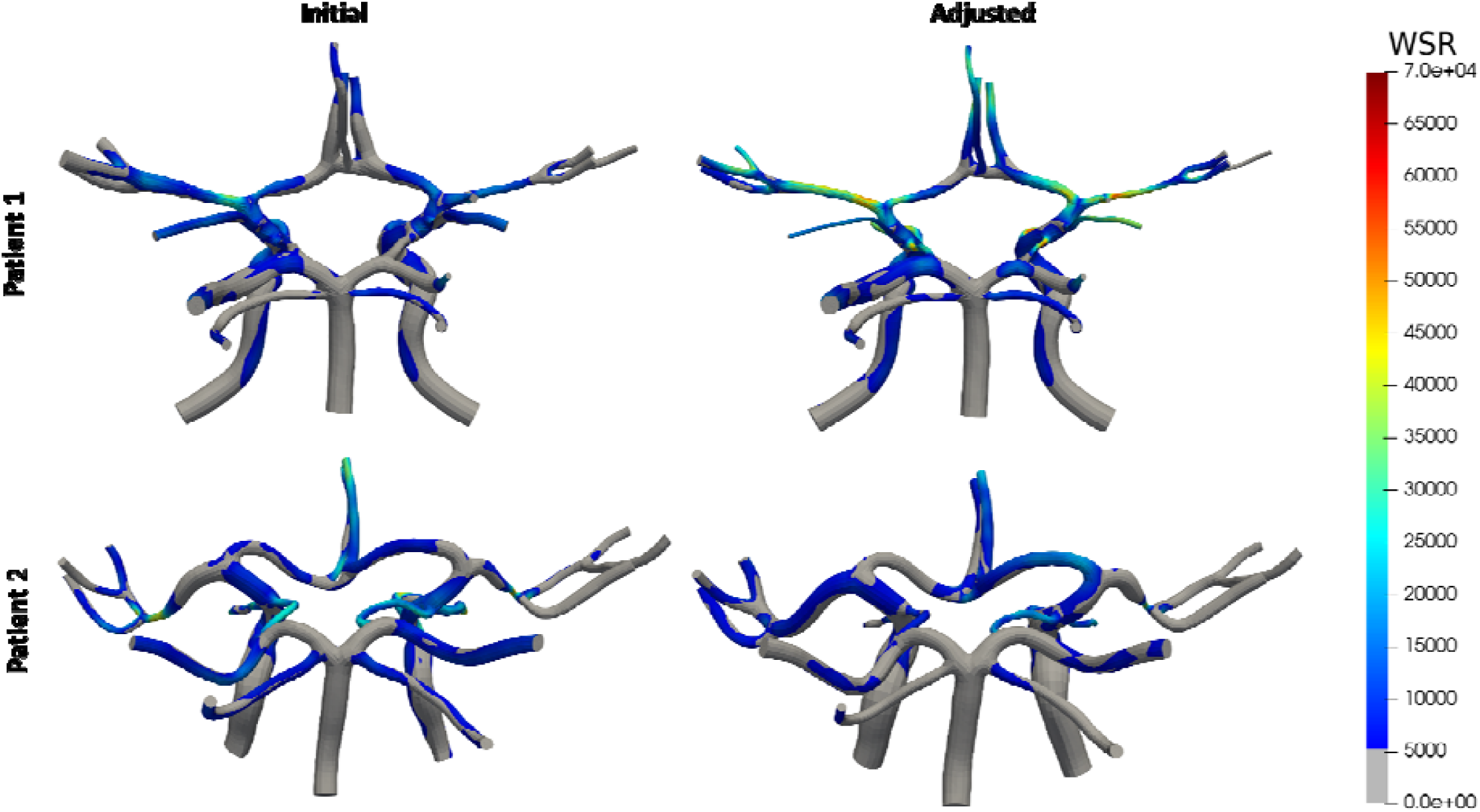
Comparison of predicted WSR (above 5000 s^-1^ coagulation limit) distributions for the initial (left panels) and XA-adjusted (right panels) models of patient 1 (top) and patient 2 (bottom).

To determine if the XA-based adjustments of the two patients’ CoW models improve model accuracy, we compared the geometries to clinical CTA data that was available for each patient. Specifically, cross-sectional areas taken from each model were compared with cross-sectional areas extracted from corresponding CTA slices, and the deviations between the two were quantified. Fig. 7 shows NRMSDs between the model cross-sectional areas and CTA lumen areas along each vessel for the initial and adjusted models for both patients. In most of the vessels considered for the two patients, the adjusted model shows less deviation from the cross-sectional areas suggested by the CTA images. Of note, the NRMSD for the RACA in patient 1 decreased from 182% (of the average CTA area in the RACA) to only 48% after adjustment. However, two of the six vessels of interest for patient 1 had more deviation from the CTA images after adjustment. The NRMSD increased from 31% to 44% for the LMCA after adjustment and it increased from 26% to 32% for the RICA. In patient 2, the RICA had increased NRMSD after adjustment (36% vs 31% in the initial model), but the remaining vessels of interest exhibited smaller NRMSD in the adjusted model vessels compared to the initial model.

**Fig 7:**
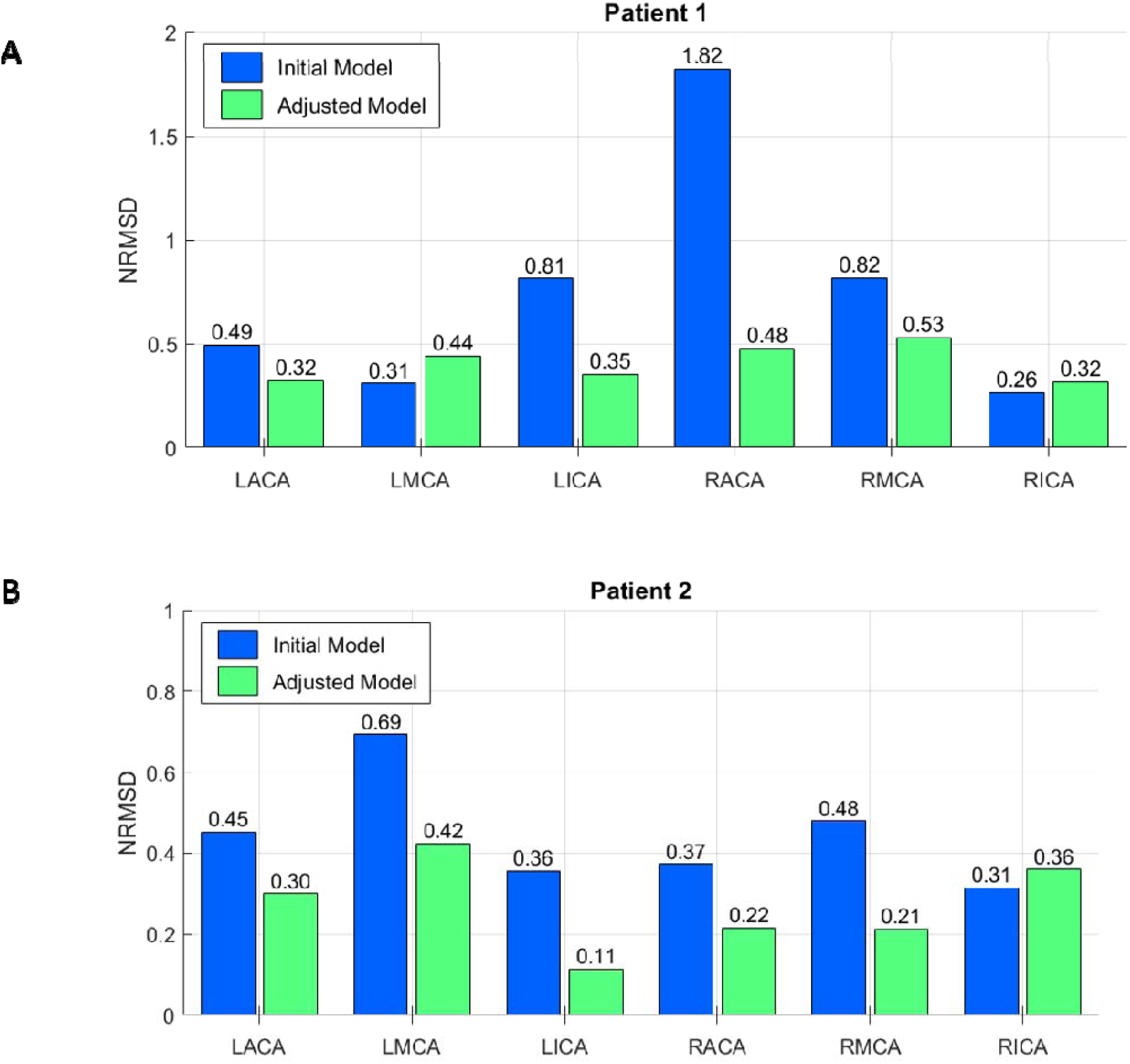
Comparison of NRMSD values for each vessel before (blue bars) and after XA-based adjustment (gold bars) for (A) patient 1 and (B) patient 2.

## 4. Discussion

With growing focus on expanding personalized medicine, patient specific geometries have been increasingly used to make personalized assessments of disease status and/or enable tailored planning of treatment strategies. Recently, we performed patient-specific hemodynamic analysis of the CoW with the goals of assessing stroke risk in pediatric MMD patients and potentially aiding clinicians in determining if surgical intervention may be required^8^. A common, and often preferred, source for patient-specific vascular geometries is clinically obtained CTA volumetric data that is segmented to create 3D vascular models. However, from a retrospective study of 50 patients we conducted in a previous work^8^, this type of imaging is not generally used in the pediatric MMD patient population as part of standard clinical follow-up practice. Instead, 3D MR TOF imaging is typically used. This imaging modality has inherent challenges that make it susceptible to insufficient image quality, which can compromise the accuracy of any 3D vascular model created from it. Complex and/or slow flow patterns are usually insufficiently resolved which can introduce inaccuracies in areas of vessel narrowing – a major clinical feature of interest in MMD. To improve the accuracy of patient specific models derived from MR TOF data, we have developed a method to locally correct vessel geometry using 2D XA imaging data that is often collected during MMD patient evaluation.

Using the XA-based adjustment process, we can generate 3D vascular models that show improved agreement with XA images, which is considered the gold standard for lumen caliber evaluation^24^. In the present study, we have adopted this approach for two patients selected based on the availability of clinical data using all three of the following imaging modalities: MR TOF, XA, and CTA. The goal is to compare the XA-adjusted CoW model with that seen in CTA images of the same patient. Given the rarity of such cases, only patient 1 is a pediatric MMD patient. Patient 2 is a non-MMD patient who presented with clinical symptoms of right sided stroke and whose imaging revealed evidence of a focal high-grade stenosis in the RMCA. When we compare patient specific geometries before and after XA-based adjustment, we see significant changes in local vessel geometry including more accurate representations, relative to XA images, of vessels that exhibit MMD-related vessel narrowing as noted by the patient’s history. For example, patient 1’s RMCA and LACA were identified by clinicians as exhibiting narrowing and both vessels are significantly impacted by the XA-adjustment process where the average vessel diameters were reduced by 48% and 30% after adjustment in the two vessels respectively. In patient 2, one of the most impacted vessels was the LMCA which increased in local diameter by 132% where the initial model was the narrowest.

We performed hemodynamic simulations on the initial and adjusted geometries of both patients and demonstrated that the distribution and magnitudes of WSR values in the CoW can be highly sensitive to geometric variations with potential implications for clinical relevance of the results. The simulation results show that, following adjustment, the surface averaged WSR in the full CoW models at peak systole increased by 86% in patient 1 (from 2.1x that of a healthy control case^8^ to 3.9x the control) and decreased by 40% in patient 2 (from 3.1x to 1.9x that of the control) relative to the initial models. Furthermore, in these two patients the maximum WSR in several vessels of interest at peak systole changed by more than 70% after adjustment. For patient 1, the presence of focal regions of critically high WSRs at least 120% greater than seen in the previously simulated control in 5 of 6 vessels of interest of the adjusted model may be an indicator of impending secondary stroke contralateral to the primary stroke event. However, the results from the initial model, with fewer areas of severe WSR (only one vessel with max WSR greater than 20% of the that seen in the control), might not indicate stroke risk at all, thus negatively impacting preventive intervention measures. This underscores the need for accurate patient specific geometries in noninvasive assessment of stroke risk in pediatric cerebrovascular disease and guiding the timing of surgical intervention.

While XA-based adjustment allows us to locally correct a 3D vascular model such that it agrees with the vessel geometry depicted on XA images, we currently do not know if the adjusted model is more representative of the patient’s 3D vessel anatomy than the initial MR-segmented model. To verify that our adjustment methods improve vascular model accuracy, we use CTA imaging as an additional reference for comparison. CTA is widely used for 3D vascular model reconstruction and provides stronger signal in the vasculature, particularly in narrowed vessels, than MRI data^25, 26^. Thus, we consider the CTA volume a more accurate 3D representation of the patient’s CoW compared to MR TOF and examine if the adjusted model more closely matches the CTA images than the initial MR-derived model. The agreement between model and CTA was assessed by comparing the cross-sectional areas of the vessels of the model and of the vessels depicted in the CTA volume. Cross-sectional areas were collected at several points throughout the vessel and a NRMSD was calculated for each vessel of the initial and adjusted models. The NMRSD for a vessel gives a measure of how much, on average, the local vessel size in the model deviates from what the CTA images suggest the vessel size should be. For all vessels of interest (in both patients) except for three, the NRMSD is reduced following adjustment. This indicates that overall, the adjusted models more closely match the CTA images than the initial models and is evidence that the XA-based adjustment procedure may improve accuracy of MR-derived vascular models.

While these results are promising and support the effectiveness of our proposed adjustment procedures, the verification approach used herein has limitations. For our purposes, the CTA volumetric data is considered as accurate representations of the patient-specific vascular anatomy. While CTA is commonly used to this end in computational studies, whether the CTA data provides good “ground truth” of target geometries is unknown. Factors including poor slice resolution, low contrast in smaller vessels, bone artifacts, and suboptimal synchronization between the contrast injection and image acquisition negatively impact the accuracy of the vessel representations in the CTA volume. Additionally, to extract the vessel cross-sectional areas from the CTA slices, we use algorithmic lumen boundary detection guided by the input of an expert pediatric neuroradiologist. These algorithms rely on image pixel intensity which can vary from vessel to vessel within the same CTA volume and among different CTA volumes. Thus, defining a standard method for identifying the lumen boundary is challenging. Furthermore, the clinician’s assessment of vessel caliber was done solely based on visual inspection where subjectivity and factors such as image brightness and contrast can impact the consistency of the results slice to slice. To minimize human error, a more comprehensive analysis should be performed utilizing inputs from multiple trained neuroradiologists.

The above challenges in assessing the accuracy of MR-derived and XA-adjusted models all originate from the need of a known target geometry for comparison. In the future, additional verification of our XA-based adjustment method using known CoW geometries is planned. The goal is to generate multiple physical 3D printed CoW models of known geometry, collecting MR TOF and XA imaging data of the physical models while mimicking the clinical settings for patient imaging, and reconstructing XA-adjusted MR-derived vascular models from the imaging data following the procedures discussed here. These models can then be directly compared to the corresponding known geometries to determine how well the XA-based adjustment performs and whether it produces a more accurate model.

In summary, patient-specific modeling of vascular networks has proven to be a valuable tool in the research of vascular pathologies. In a previous work, we showed how patient-specific hemodynamic analysis of the CoW and WSR can be used to noninvasively assess stroke risk in pediatric MMD, potentially resulting in earlier diagnosis of disease progression with reduced stroke burden and improved clinical outcome. These analyses rely on geometrically authentic 3D vascular models that can be constructed using volumetric CTA or MR imaging data, the latter of which is the more likely imaging modality for pediatric MMD patients due to radiation exposure concerns with CTA. However, MR imaging can be suboptimal for 3D model reconstruction because of insufficient image resolution. We developed a method where virtual angiographies of 3D MR-derived vascular models are conducted and the resulting 2D projections are compared to corresponding 2D XA images, which are commonly used to clinically assess vessel narrowing. The 3D model is then locally adjusted until its 2D projections and XA images are in agreement. Through blood flow simulations, we demonstrated that local WSR distributions in an initial MR-derived model is considerably different from the XA-adjusted model which can lead to incorrect stroke risk assessments. A comparative analysis with patients’ CTA imaging data suggests that XA-based adjustment can improve vascular model accuracy and enable more accurate assessments of stroke risk, thereby improving individualized treatment and monitoring of pediatric MMD patients.

## Acknowledgements

The authors gratefully acknowledge Texas Advanced Computing Center (TACC) for providing high-performance computing resources.

## Sources of funding

This work was supported by NIH grant R03NS110442 to SSH.

## Conflict of interest/Disclosure

ZS is a stockholder in Alzeca Biosciences and a consultant for InContext.ia. All other authors declare that they have no conflict of interest.

## Ethical approval

There was no direct involvement of human subjects or protected health information (PHI). All patient imaging data used in analysis and modeling was collected retrospectively from medical records and de-identified at Texas Children’s Hospital. Institutional review board (IRB) approval was obtained with a waiver of written authorization for consent.

